# The FDA-approved CDK4/6 inhibitor Abemaciclib is a potent agonist of the β-arrestin-biased atypical chemokine receptor ACKR3

**DOI:** 10.64898/2026.11.03.764001

**Authors:** Maurice C.M.L. Buzink, Joy Darcis, Desislava Nesheva, Max Meyrath, Shawn Jenjak, Nilanjana Banerjee, Manisankar Ganguly, Ashna Reyaz, Ramanuj Banerjee, Esther A. Leliveld, Yong-Jun Kwon, Jerome Paggetti, Sybrand Zielhuis, Dick Pluim, Alwin Huitema, Brian F. Volkman, Henry F. Vischer, Arun K. Shukla, Andy Chevigné, Martyna Szpakowska, Rob Leurs

## Abstract

Atypical chemokine receptor 3 (ACKR3/CXCR7) is an intrinsically β-arrestin-biased family-A GPCR that controls chemokine availability by scavenging CXCL12 and CXCL11, and consequently plays a key role in cancer, chronic inflammation, autoimmunity, cardiovascular disease, and CNS repair. Despite its pathophysiological significance, the chemical space available to interrogate and therapeutically modulate this receptor remains limited. Here, by using a drug-repurposing strategy to screen 2,342 FDA-approved compounds in a NanoBRET-based fluorescent chemokine-binding competition assay, we identified the clinically used CDK4/6 inhibitor abemaciclib (Verzenios®) as a nanomolar ACKR3 ligand. Abemaciclib inhibited CXCL12 binding to ACKR3 with comparable affinity to the ACKR3-reference agonist VUF15485, whereas the related clinically used CDK4/6 inhibitors palbociclib, ribociclib, dalpiciclib and trilaciclib were inactive on this receptor. Binding and functional profiling across classical and atypical chemokine receptors revealed selectivity for ACKR3, including lack of detectable activity at the CXCL12 receptors CXCR4 and ACKR5. Abemaciclib acted as a potent ACKR3 agonist, recruiting both β-arrestin-1 and 2 and promoting rapid receptor internalization with potencies and efficacies, like the reference ACKR3 agonist VUF15485. Major CYP3A4-derived abemaciclib metabolites retained ACKR3 binding and agonist activity, albeit with reduced potency. Methionine ¹³C-HSQC NMR confirmed direct receptor engagement and revealed a small-molecule-like ACKR3 conformational signature distinct from the CXCL12-bound state. Cryo-EM of the abemaciclib–ACKR3–CID24 complex at 3.2 Å uncovered a unique, inverted U-shaped ligand binding pose in the orthosteric pocket and activation-associated rearrangements of TM6, TM7 and helix 8. Functionally, abemaciclib blocked ACKR3-mediated uptake of CXCL11 and CXCL12 *in vitro*. Moreover, oral administration of abemaciclib in mice resulted in a time-dependent increase in plasma CXCL12 levels, indicative of *in vivo* ACKR3 target engagement. These findings establish this FDA-approved kinase inhibitor as a potent ACKR3 modulator and reveal an unexpected pharmacological link of this CDK4/6 inhibitor to atypical chemokine receptor biology.

## Introduction

Atypical chemokine receptor 3 (ACKR3), also known as CXCR7, is the most studied member of the atypical chemokine receptor family and a prototypical β-arrestin-biased G protein-coupled receptor (GPCR)(1–3). ACKR3 was initially deorphanized as a high-affinity chemokine receptor for CXCL12 (4) and subsequently CXCL11 (5), chemokines long considered to signal exclusively through CXCR4 and CXCR3, respectively. Although ACKR3 retains the canonical seven-transmembrane architecture of class A GPCRs, it generally lacks detectable G protein activation and instead preferentially recruits GPCR kinases (GRKs), β-arrestins and receptor trafficking machinery both constitutively and upon agonist stimulation (6–11). The lack of canonical G protein-mediated signalling but observed internalization early on led to the suggestion that ACKR3 acts as a “chemokine sink” or as a “decoy receptor”, and thereby shapes chemokine availability, gradient formation, and receptor responsiveness in tissue microenvironments (12–14). Beyond chemokines, ACKR3 has recently also been shown to bind and scavenge diverse endogenous peptides, including the proadrenomedullin N-terminal 20 peptide 12 (PAMP-12) and opioid peptides such as enkephalins, dynorphins, and BAM22, suggesting a broad role in regulating extracellular signalling events (2,15,16). Moreover, with the recognition of β-arrestin-mediated intracellular signalling (17), the roles of the β-arrestin biased atypical chemokine receptors in cellular communication have become even more intriguing (3).

Despite the lack of detailed understanding of the role of ACKRs in cellular signalling, their translational potential has already been described for several important therapeutic areas. ACKRs are suggested to reshape pathological chemokine networks in cancer, chronic inflammation, autoimmunity, cardiovascular disease and CNS repair (1,3,18,19), with most of the current attention directed to ACKR3. The therapeutic implications of ACKR3 are broad but context dependent. In cancer, ACKR3 is overexpressed in multiple tumour types, including breast, lung and brain cancers, and has been associated with proliferation, migration, angiogenesis, metastasis and therapy resistance (20). In demyelinating disease, ACKR3 antagonism has generated particular interest. In the adult central nervous system, CXCL12– CXCR4 signalling supports oligodendrocyte precursor-cell maturation and remyelination, whereas ACKR3-mediated CXCL12 scavenging can restrict local CXCL12 availability (21). Pharmacological ACKR3 antagonism enhanced oligodendrocyte maturation and myelin repair in preclinical demyelination models (21,22). These data support a therapeutic concept in which ACKR3 blockade simultaneously modulates neuroinflammation and enhances repair by increasing accessible CXCL12 in relevant CNS niches. Finally, in cardiovascular disease, ACKR3 has been linked to endothelial biology, platelet activation, atherosclerosis, myocardial injury, pain modulation, and tissue repair (19,23,24).

To further delineate the translational potential of ACKR3 targeting, there is a strong need to identify chemically diverse sets of small-molecule scaffolds targeting ACKR3. Despite the availability of several useful pharmacological tools, including ACKR3 agonists such as CCX662 (25), VUF15485 (26), LIH383 (16) and conolidine (24) and the antagonists ACT-1004-1239 (22,27–29), and VUF16840 (9), the current chemical space remains narrow and is dominated by a limited number of chemotypes (30,31). Structurally diverse ligands would also support structure–activity and structure–function studies, helping to define whether ACKR3 can be selectively biased toward therapeutically beneficial outcomes.

In search of new chemical entities targeting ACKR3, we employed a drug repurposing strategy. Drug repurposing can accelerate ACKR3 therapeutic exploration by starting from compounds with known pharmacokinetic and safety profiles, thereby reducing early development risks (32,33). As a result of screening a library of FDA-approved drugs, we identified abemaciclib (Verzenios® - a clinically used cyclin-dependent kinase 4 and 6 inhibitor) as a new nanomolar affinity ligand for ACKR3. Moreover, abemaciclib exhibits excellent selectivity for ACKR3 and does not interact with other (atypical) chemokine receptors. Using ^13^C-Methionine ACKR3 protein NMR and cryo-EM studies, detailed information on the interaction of the kinase inhibitor with ACKR3 was obtained. Functionally, abemaciclib affected ACKR3-mediated chemokine uptake both *in vitro* and *in vivo*.

## Results

### The CDK4/6 inhibitor abemaciclib (Verzenios®) is a high-affinity, selective ACKR3 binder

In search for new chemical scaffolds for ACKR3 ligands we employed a repurposing approach and screened the FDA-approved drug library from Targetmol for ACKR3 binders. We utilized a NanoBRET-based fluorescent chemokine binding assay, using 10 µM of each of the 2342 displacers, 0.3 nM fluorescently labelled CXCL12-AF647 and HEK-293T membranes, expressing human ACKR3, N-terminally tagged with Nanoluciferase (NLuc) (**Fig. 1A**). In this screen, we identified the CDK4/6 inhibitor abemaciclib as a full inhibitor of CXCL12-AF647 binding to ACKR3 (**Fig. S1**). To validate the initial screening results, fresh compound solutions were prepared and NanoBRET competition binding was repeated with a full range of concentrations, also including the reference ACKR3 ligand VUF15485 (26). Surprisingly, abemaciclib effectively displaced the binding of CXCL12-AF647 to ACKR3 in nanomolar concentrations resulting in a pK_i_ value of 7.6 ± 0.4 (mean ± SD, n = 3) and an affinity value of 25 nM. In the same assay a similar nanomolar affinity for ACKR3 was obtained for the reference agonist VUF15485 (pK_i_ = 7.8 ± 0.2, n = 3).

**Figure 1:**
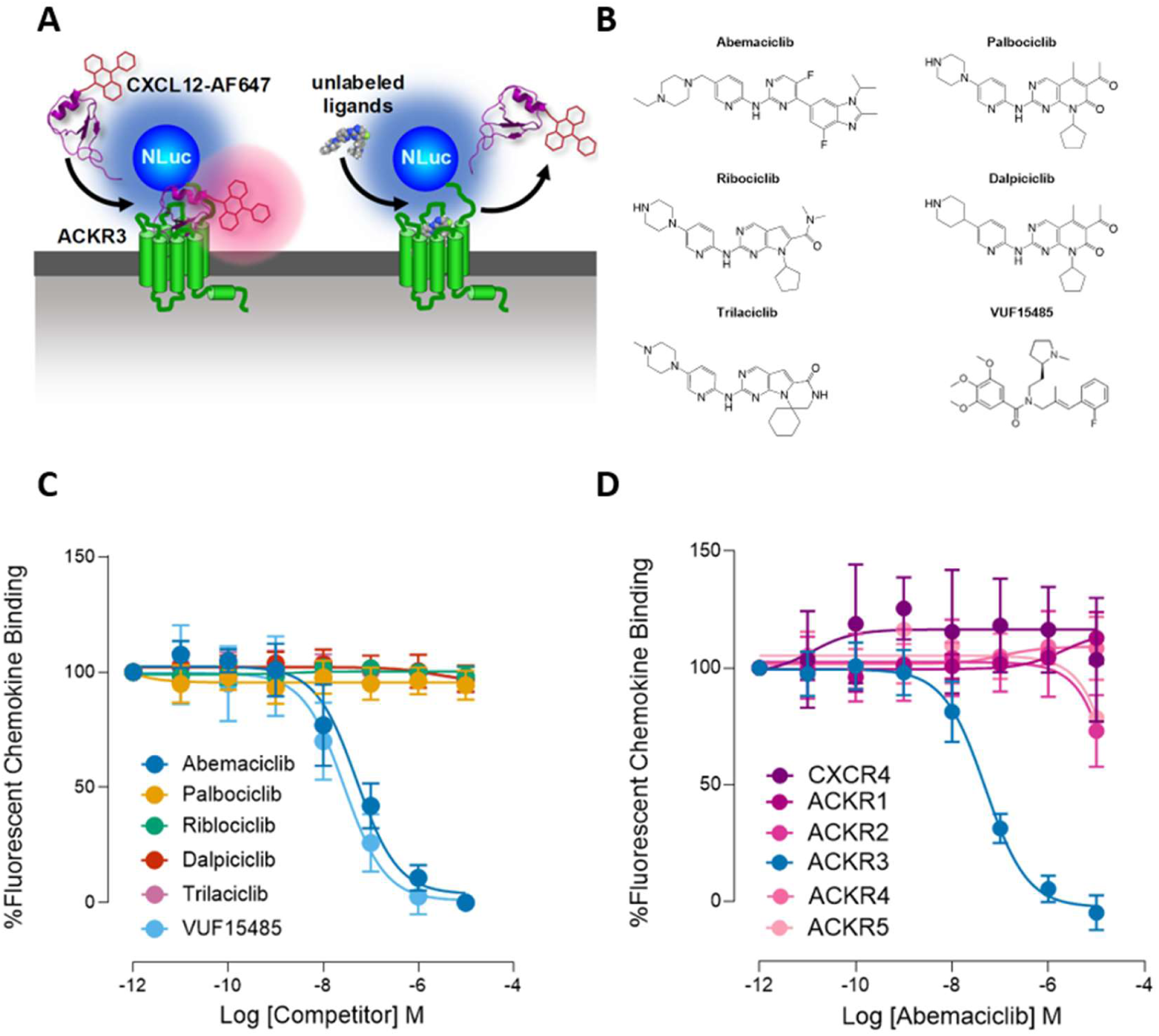
Abemaciclib binds to ACKR3 in a NanoBRET-based chemokine binding assay. A) Schematic representation of the employed NanoBRET-based binding assay, using fluorescently labelled CXCL12-AF647 and ACKR3 with the bioluminescent enzyme Nluc fused to its N-terminus. B) Structures of abemaciclib, reference agonist VUF15485 and clinically approved CDK4/6 inhibitors C) Membranes of HEK-293T cells transiently expressing NLuc-ACKR3 were used to measure the concentration dependent displacement of binding of 0.3 nM CXCL12-AF647. Percentage of remaining CXCL12-AF647 binding was normalized. D) Membranes of HEK-293T cells transiently expressing either NLuc-ACKR1, NLuc-ACKR2, NLuc-ACKR3, NLuc-ACKR4, NLuc-ACKR5 or NLuc-CXCR4 were used to measure the concentration dependent displacement of binding of 10 nM CXCL11-AZ594, 10 nM CCL5-AZ594, 0.3 nM CXCL12-AF647, 10 nM CCL19-AZ594, 10 nM CXCL12-AZ488 or 10 nM CXCL12-AF647, respectively. Data was normalized to percentage of remaining fluorescently labelled chemokine compared to vehicle. Graphs represent pooled data from 3 independent experiments. Data points depicted are means, the error bars represent respective S.D. values.

Abemaciclib is an FDA-approved dual inhibitor of cyclin-dependent kinase CDK4 and CDK6. Commercially known as Verzenios®, abemaciclib is successfully used in the clinic for the treatment of HR-positive, HER2-negative, lymph node-positive, high-risk early breast cancer (34). Next to abemaciclib, the related CDK4/6 inhibitors palbociclib, ribociclib, dalpiciclib, and trilaciclib (see **Fig. 1B** for structures) have been globally approved for clinical use (35). Therefore, as the next step, we tested all clinically approved CDK4/6 inhibitors for their ability to interact with ACKR3. As can be seen in **Fig. 1C**, none of the other CDK4/6 inhibitors affected the binding of CXCL12-AF647 to the NLuc-tagged ACKR3 i.e., the observed NanoBRET signal remained approximately 100%.

Since CXCL12 does not only bind ACKR3, but is also the endogenous ligand for the chemokine receptor CXCR4 and the atypical chemokine receptor ACKR5 (2,36–38), formerly known as GPR182, we tested if abemaciclib displays affinity for these targets too. However, in NanoBRET-based fluorescently labelled CXCL12 binding assays for Nluc-tagged CXCR4 and ACKR5, abemaciclib showed no affinity for either of these other classical and atypical receptors (**Fig. 1D**). Specifically, abemaciclib showed no fluorescent tracer displacement from CXCR4 or ACKR5 unlike IT1t or unlabelled CXCL12 respectively (**Fig. S2**). Moreover, abemaciclib did not show affinity for any of the other atypical chemokine receptors, (**Fig. 1D**, **Fig. S2**). Finally, ACKR3 binds not only CXCL12 but also several other endogenously occurring chemokines, including CXCL11 (2). As can be seen in **Fig. S3**, abemaciclib also effectively inhibited binding of fluorescent CXCL11-AZ594 to NLuc-tagged ACKR3.

### Abemaciclib increases β-arrestin recruitment to ACKR3 and its internalization

The atypical chemokine receptor ACKR3 is a biased chemokine receptor, which does not signal via canonical G-protein pathways but can recruit β-arrestins following agonist stimulation (8,39). To evaluate the effect of abemaciclib binding to ACKR3, we tested abemaciclib together with reference ACKR3 agonist VUF15485 and inverse agonist VUF25550 (also known as ACT-1004-1239) in a NanoBiT-based β-arrestin-2 recruitment assay in transiently transfected HEK-293T cells (**Fig. 2A**). As can be seen, both abemaciclib (1 µM) and VUF15485 (1 µM) rapidly induced a strong β-arrestin-2 recruitment (**Fig. 2C**). In line with earlier findings (9), constitutive β-arrestin-2 recruitment by ACKR3 was observed in the present assay. This basal ACKR3-β-arrestin-2 recruitment was rapidly inhibited by the inverse agonist VUF25550 (**Fig. 2C**). The observed effects were concentration-dependent (**Fig. 2D**) with pEC_50_ values of 7.8 ± 0.4 and 8.1 ± 0.2 (mean ± SD, n = 3) for the agonists abemaciclib and VUF15485, respectively. In the same assay, the inverse agonist VUF25550 potently inhibited the constitutive ACKR3 activity with a pIC_50_ value of 9.8 ± 0.7 (mean ± SD, n = 3).

**Figure 2:**
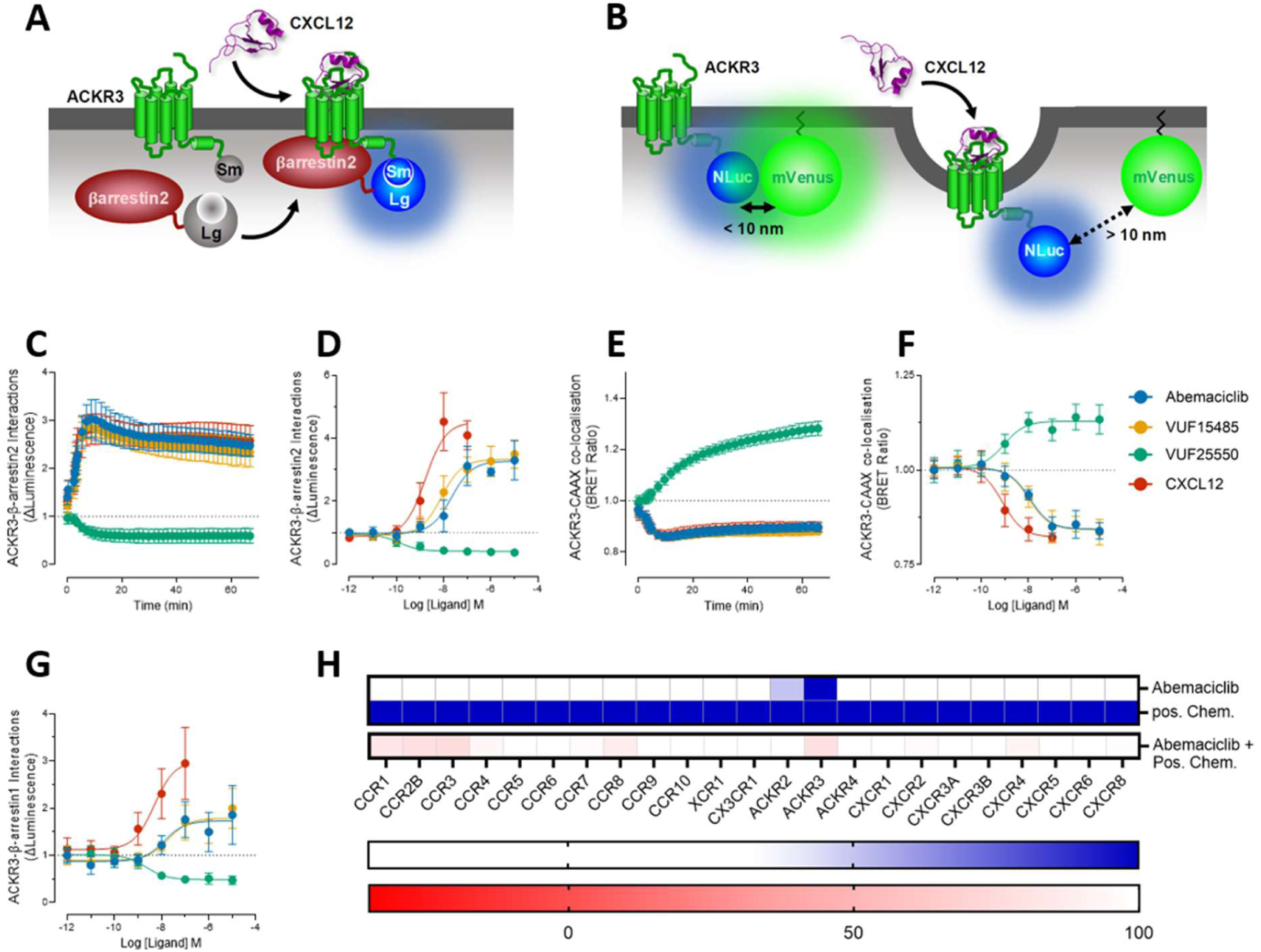
Abemaciclib is a selective agonist for ACKR3. A) Schematic representation of the NanoBiT complementation technology to detect interactions between ACKR3-SmBiT and LgBiT-ß-arrestin2. B) Schematic representation of the BRET system to detect ACKR3-NLuc interactions with mVenus-CAAX. C) HEK-293T cells were transfected with ACKR3-SmBiT and LgBiT-ß-arrestin2 and measured over time with 1 µM of abemaciclib, VUF15485, VUF25550, or CXCL12. D) HEK-293T cells were transfected with ACKR3-SmBiT and LgBiT-ß-arrestin2 and stimulated for 15 minutes with abemaciclib, VUF15485, VUF25550, or CXCL12. Recruitment of ß-arrestin2 to ACKR3 is visualized as luminescence fold over vehicle. E) HEK-293T cells transfected with ACKR3-NLuc and mVenus-CAAX were used to measure the ligand-induced ACKR3 co-localization with CAAX over time after the addition of 1 µM of abemaciclib, VUF15485, VUF25550, or CXCL12. F) HEK-293T cells were transfected with ACKR3-NLuc and mVenus-CAAX and stimulated for 15 minutes with abemaciclib, VUF15485, VUF25550, or CXCL12. G) HEK-293T cells were transfected with ACKR3-SmBit and LgBit-ß-arrestin1 and stimulated for 15 minutes with abemaciclib, VUF15485, VUF25550, or CXCL12. Recruitment of ß-arrestin1 to ACKR3 is visualized as luminescence fold over vehicle. C-G) Graphs represent pooled data from 3 independent experiments. Data points depicted are means, the error bars represent respective S.D. values. H) HEK-293T cells were transfected with vectors encoding chemokine receptors C-terminally fused to SmBiT and LgBiT-ß-arrestin2. The recruitment induced by 10 µM abemaciclib (10 µM) or positive control chemokine (200 nM) was measured for 20 minutes after stimulation of each chemokine receptor (Blue). Antagonist effect of abemaciclib was assessed in the presence of full agonist chemokine for each receptor (Red). Data are presented as percentage of full agonist response.

As ACKR3 is known to recruit both β-arrestin-1 and β-arrestin-2 (40,41), abemaciclib was tested in an ACKR3-mediated β-arrestin-1 recruitment assay (**Fig. 2G**). In a similar NanoBiT-based assay configuration, both the reference agonist VUF15485 and abemaciclib concentration-dependently recruited β-arrestin-1 to ACKR3 with pEC_50_ values of 7.7 ± 0.2 and 7.9 ± 0.3 (mean ± SD, n = 3), respectively. This assay included the reference VUF25550 ligand which acted as inverse agonist (**Fig. 2G**, pIC_50_ = 8.7 ± 0.3, mean ± SD, n = 3).

ACKR3 is also known to rapidly internalize, both constitutively and upon agonist exposure (9,42). To measure ACKR3 internalization, the receptor protein was C-terminally tagged with nanoluciferase and transiently co-expressed in HEK-293T cells alongside membrane-localized mVenus-CAAX (**Fig. 2B**). Co-localization of ACKR3 and mVenus-CAAX in the plasma membrane results in a NanoBRET signal, which is reduced upon receptor internalization. As observed in the β-arrestin-1 and -2 recruitment assays, the ACKR3 agonists VUF15485 and abemaciclib increased ACKR3 internalization within minutes, manifested as a reduction in the NanoBRET signal (**Fig. 2E**). Interestingly, exposure to the inverse agonist VUF25550 resulted in a robust increase of the NanoBRET signal, indicative of an increased membrane expression of ACKR3 upon inhibition of constitutive ACKR3 internalization, as also previously reported (9,42). Again, all the observed effects on ACKR3 membrane translocation were concentration-dependent (**Fig. 2F**), yielding pEC_50_ values of 7.9 ± 0.1 and 8.0 ± 0.2 (mean ± SD, n ≥ 3) for the agonists abemaciclib and VUF15485, respectively, and a pIC_50_ value of 9.1 ± 0.2 (mean ± SD, n = 3) for the inverse agonist VUF25550.

### Abemaciclib selectively activates ACKR3

As shown in **Fig. 1D**, the results from the NanoBRET-based fluorescent chemokine binding assays for ACKR1-ACKR5 indicate that abemaciclib is a selective ACKR3 ligand. To further substantiate this, abemaciclib was functionally tested in agonist-and antagonist modes against the complete panel of chemokine receptors. The human chemokine receptor family comprises 23 GPCRs: 18 “classical” G-protein signalling receptors grouped by ligand class into CCR, CXCR, CX3CR and XCR subfamilies, plus the five atypical chemokine receptors (2). To test the selectivity of abemaciclib within the chemokine receptor family, a NanoBiT-based β-arrestin-recruitment assay was employed for the large panel of chemokine receptors, as described (16). As **Fig. 2H** shows, ACKR3 was the only chemokine receptor activated by abemaciclib (10 µM). Moreover, chemokine-induced β-arrestin2 recruitment to none of the other chemokine receptors was seriously affected by 10 µM abemaciclib when tested in antagonist-mode. These findings are in line with the earlier data obtained in the NanoBRET-based binding assays in **Fig. 1D**.

### Abemaciclib metabolites effectively bind and activate ACKR3

Abemaciclib is extensively metabolized in the liver primarily by the cytochrome P450 enzyme CYP3A4 into several active metabolites, the most notable being N-desethyl-abemaciclib (M2), hydroxyabemaciclib (M20) and hydroxy-N-desethylabemaciclib (M18) (**Fig. 3A**) (43–45). As can be seen in Fig. 3, the three main abemaciclib metabolites were also able to bind (**Fig. 3B**) and activate (**Fig. 3C**) ACKR3, with the M18 metabolite showing the largest reduction in affinity and consequently, potency.

**Figure 3:**
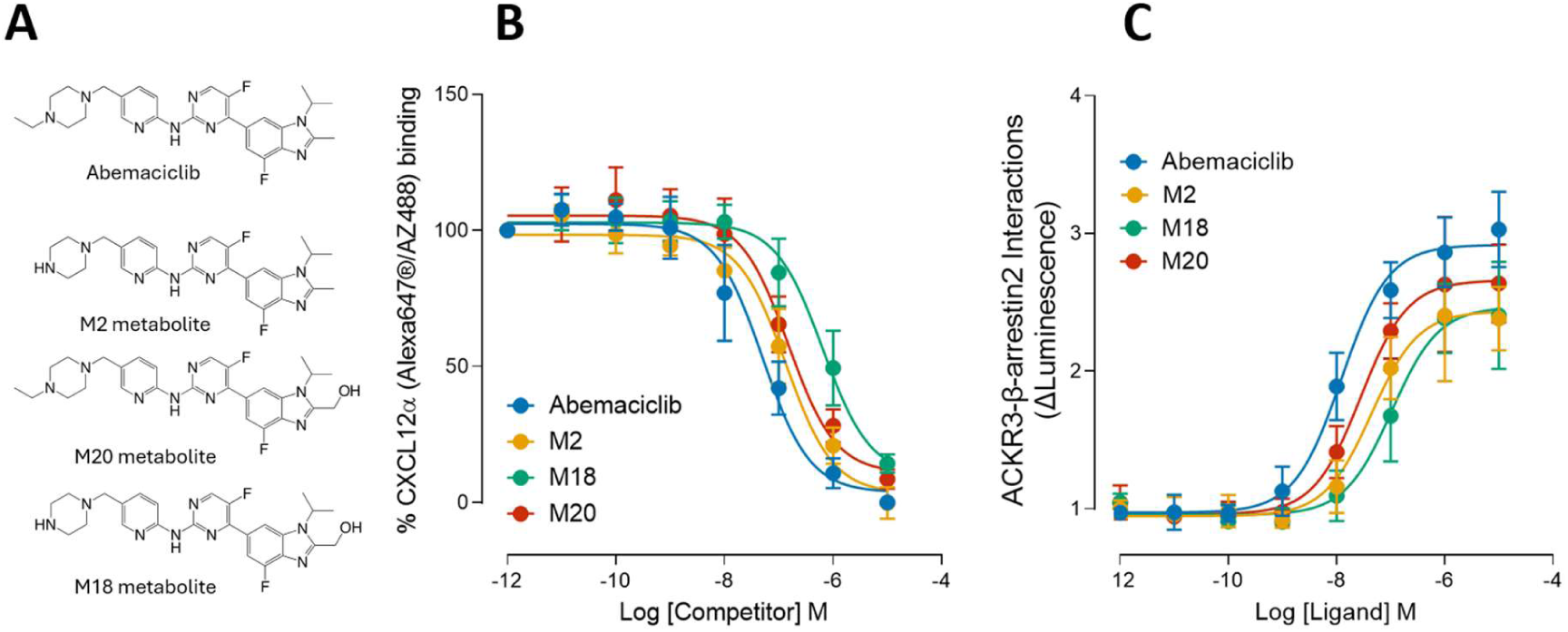
Abemaciclib metabolites bind and activate ACKR3. A) Structures of abemaciclib and abemaciclib metabolites M2, M18, and M20 B) Membranes of HEK-293T cells transiently expressing NLuc-ACKR3 were used to measure the concentration dependent displacement of binding of 0.3 nM CXCL12-AF647. Percentage of remaining CXCL12-AF647 binding was normalized. Obtained pIC50 values were 7.4 ± 0.4, 6.9 ± 0.3, 6.2 ± 0.1 and 6.8 ± 0.2 (mean ± SD, n=3) for abemaciclib, M2, M18, and M20, respectively. C) HEK-293T cells were transfected with ACKR3-SmBit and LgBit-β-arrestin2 and stimulated for 15 minutes with abemaciclib or metabolites M2, M18, or M20. Recruitment of β-arrestin2 to ACKR3 is visualized as luminescence fold over vehicle. Obtained pEC50 values were 7.9 ± 0.3, 7.3 ± 0.2, 7.0 ± 0.2 and 7.5 ± 0.2 (mean ± SD, n=3) for abemaciclib, M2, M18, and M20, respectively. The graphs represent pooled data from 3 independent experiments. Data points depicted are means; the error bars represent respective S.D. values.

### Abemaciclib induces a ligand-specific ACKR3 conformation

To confirm whether abemaciclib directly engages ACKR3, we compared 2D ^1^H–^13^C HSQC spectra of methionine-labeled receptor protein in several ligand-bound states (**Fig. 4A**). To evaluate these changes, we examined previously established methionine reporters mapped onto the ACKR3 structure (**Fig. 4B**). Among these, M138^3×46^, located near the conserved NPxxY motif, served as a robust and reproducible indicator of ligand-dependent conformational changes, consistent with prior work (39,46). In contrast, M212^5×39^ exhibited more variable behavior across conditions and was therefore a less reliable reporter of receptor state. Consistent with this distinction, small molecule reference ligands VUF16840 (ACKR3 inverse agonist (9) and VUF15485 (ACKR3 agonist (26) produced chemical shift changes that align with the expected movement of Met138^3×46^, whereas abemaciclib did not display clear correspondence to these canonical peak positions, instead showing less well-defined perturbations. Yet, peak shifts in the main cluster of peaks and at M138^3×46^ were clearly observed as compared to Apo-ACKR3 when abemaciclib binds ACKR3 (**Fig. 4C**).

**Figure 4:**
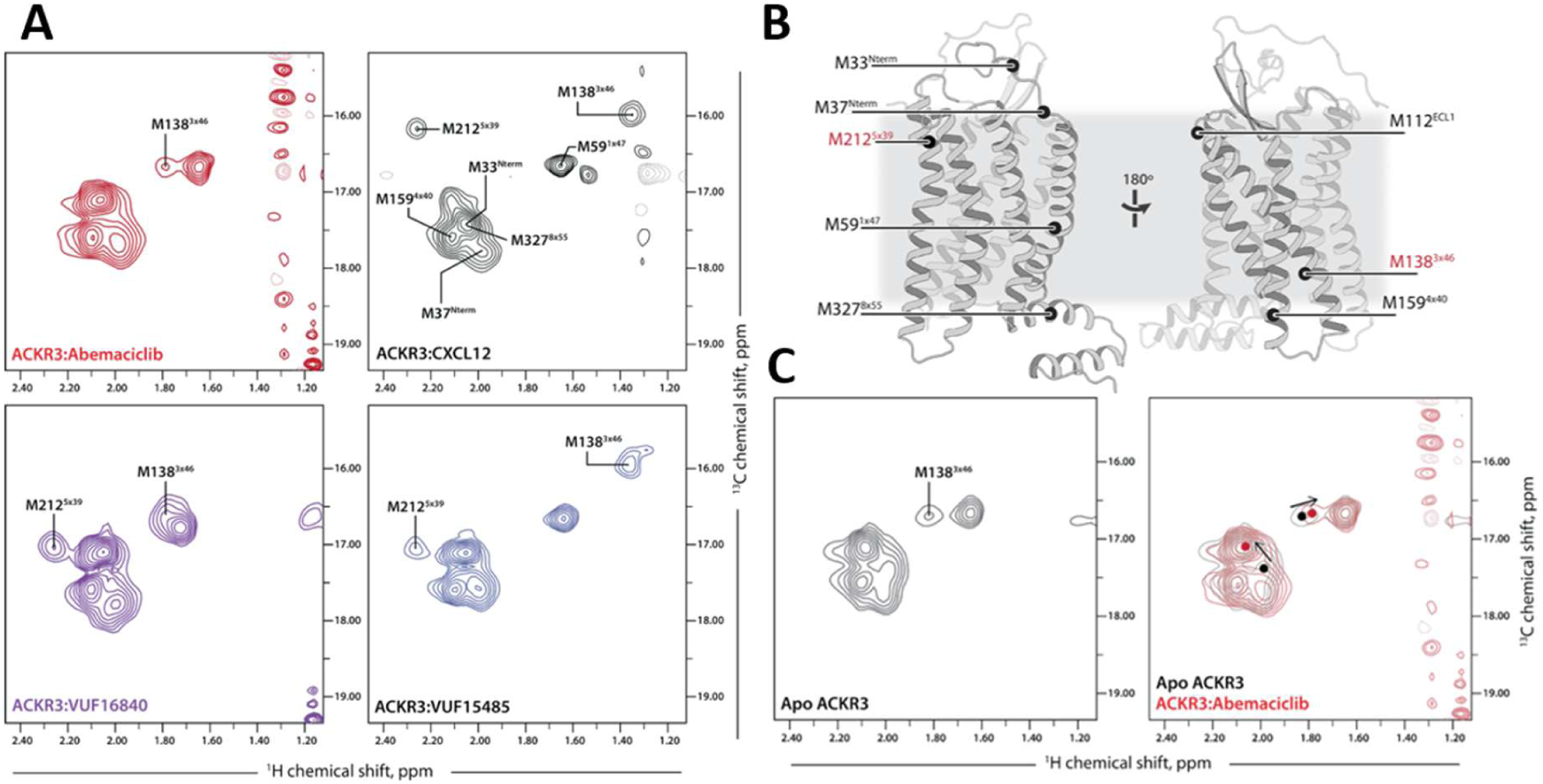
Binding of ACKR3 by abemaciclib observed by methionine 2D HSQC NMR. (A) 2D ^1^H–^13^C HSQC spectra of ACKR3 bound to several ligands at 310K. Assigned methionine residues are labeled. Negative contour peaks are semitransparent dashed lines. Abemaciclib in red, CXCL12 in black, VUF16840 in purple, and VUF15485 in blue. (B) ACKR3 with methionine residues labeled and residues of interest in red. (C) 2D ^1^H–^13^C HSQC spectra of Apo ACKR3 and an overlay with abemaciclib-bound ACKR3 with arrows to highlight peak shifts in the bound state.

Despite the ambiguity and the position of Met138^3×46^ in the abemaciclib-bound spectra of ACKR3, a cluster of methionine signals at ∼2.1 ppm (^1^H) and ∼17.5 ppm (^13^C), arising from contributions of N-terminal and transmembrane residues (including TM1 and TM4), exhibited consistent ligand-dependent behavior. Notably, the small molecule ligands VUF16840, VUF15485, and abemaciclib produced a convergent spectral pattern in this region that is clearly distinct from the CXCL12-bound state. This suggests that the small molecules stabilize a shared conformational ensemble of ACKR3 that differs from the chemokine agonist-bound conformation.

### Structural determination of the abemaciclib-ACKR3 complex using cryo-EM

To investigate the mode of binding and mechanism of ACKR3 activation by abemaciclib, we determined the structure of abemaciclib-bound ACKR3 in complex with the stabilizing antibody Fab fragment, CID24 (47) at a global resolution of 3.2 Å (**Fig 5A and S4**). The overall conformation of the abemaciclib-ACKR3 was similar to the active state structures of recently reported agonist-bound ACKR3 (47,48) and varied with a RMSD of ∼0.5 Å across the protein residues (**Fig. 5B**).

**Figure 5:**
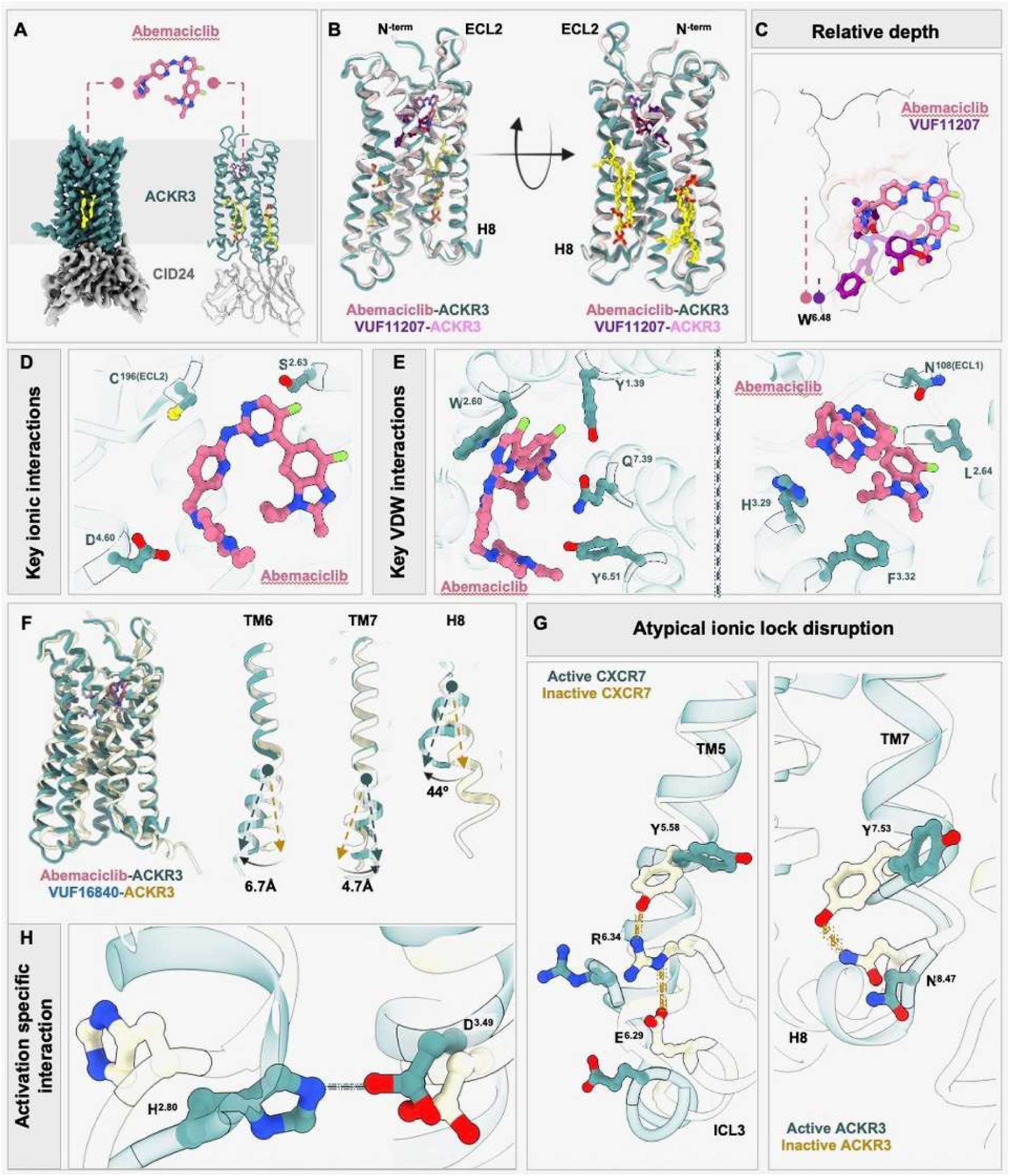
Cryo-EM structure and key features of abemaciclib bound ACKR3. A) Overall structure of abemaciclib-bound ACKR3 in complex with CID24 (PDB ID: 27SF). B) Structural alignment of abemaciclib-ACKR3 on the active state structure of VUF11207-ACKR3 (PDB ID: 9WLE). Two putative lipids bound to the TMs are indicated in yellow. C) Relative depth of insertion of abemaciclib as compared to VUF11207 within the orthosteric pocket of ACKR3 as measured from the highly conserved W^6.48^. D, E) Ball and stick representation illustrating the ionic/polar (D) and Van der Waals’ (E) interactions of abemaciclib with ACKR3. F) Superimposition of agonist abemaciclib-and inverse agonist VUF16840-bound ACKR3 structures (PDB ID: 9WLG) highlighting the movements of TM segments and helix 8 G) Disruption of the atypical ionic lock upon agonist-induced activation of ACKR3. H) Diagram highlighting the activation-specific interactions observed in the abemaciclib-ACKR3 structure.

Amino acid residues preceding Cys34^N-term^ and following Ala329^Helix-8^, corresponding to the N- and C-termini of ACKR3, were excluded from the final model because their electron densities were insufficiently defined, likely reflecting high flexibility and lack of stabilizing interactions in these regions.

Significant differences were observed in the binding mode of abemaciclib within the orthosteric pocket of ACKR3 compared to other agonist-bound ACKR3 structures. Abemaciclib adopts an inverted U-shaped conformation, harbors a buried surface area of ∼933.5 Å^2^, and docks at a depth of ∼5 Å from the conserved Trp265^6.48^ within the amphipathic orthosteric pocket. This configuration signified a shallow binding mode for abemaciclib relative to other small molecule agonists, like CXC-662 and VUF11207 (47,48), which penetrate deeper into the receptor pocket (**Fig. 5C and S4**). A total of 2 hydrogen bonds and 24 non-bonded contacts stabilize abemaciclib’s conformation within ACKR3’s orthosteric pocket. Key interactions include hydrogen bonds with Ser103^2.63^ and Cys196^ECL2^; ionic interaction with Asp179^4.60^; and additional non-bonded contacts involving Tyr51^1.39^, Asn108^ECL1^, Trp100^2.60^, Leu104^2.64^, His121^3.29^, and Phe124^3.32^, Tyr268^6.51^, and Gln301^7.39^ (**Fig 5D and 5E**).

Structural superimposition of abemaciclib bound ACKR3 with the previously reported structure of VUF16840 bound inactive structure of ACKR3 (48) revealed the hallmark conformational changes indicative of receptor activation: an outward movement of TM6 by ∼6.7 Å (as measured from the Cα of Arg251^6.34^), an inward movement of TM7 by 4.7 Å (as measured from the Cα of Ser316^7.54^), and a rotation of helix-8 by ∼44° away from the receptor (**Fig. 5F**). Local conformational variations were also observed in the N-terminus, ECL2, ICL1, and ICL3 of CXCR7 (**Fig. 5F, S4**). Additionally, local rearrangements in the rotameric conformations of residues surrounding the ligand, namely Trp100^2.60^, His121^3.29^, and Asp179^4.60^ can be observed, which appear to arise from distinct ligand conformations and docking poses, rather than from receptor activation.

Unlike most class A GPCRs, inactive ACKR3 lacks the canonical DRY-motif ionic lock, in which Arg^3.50^, Asp/Glu^3.49^, and Asp/Glu^6.30^ cooperate to stabilize the inactive conformation (48,49) In ACKR3, Glu^6.30^ is shifted to position 6.29, creating a one-residue offset in the primary sequence. In the inactive state, Arg142^3.50^ points toward the receptor core, whereas Glu246^6.29^ is oriented away from the core of the receptor. Glu246^6.29^ participates in an intrahelical ionic lock by forming a salt bridge with Arg251^6.34^, which in turn engages Tyr232^5.58^ through an interhelical ionic interaction (**Fig. 5G**). Upon activation of ACKR3 by abemaciclib, Tyr232^5.58^ rotates upwards towards the receptor core, while Arg251^6.34^ and Glu246^6.29^ shift outwards, consistent with the outward displacement of TM6 during activation (**Fig. 5G**), thereby disrupting this ionic lock.

Additionally, in the inactive state, Tyr315^7.53^ of the NPxxY motif in TM7 points downward and engages in hydrogen bonding with His80^2.40^ in TM2 and Asn319^8.47^, to limit helix 8 movement. Agonist binding triggers an upward rotation and inward shift of Tyr315^7.53^, resulting from an inward movement of TM7, that disrupts its interaction with His80^2.40^. Subsequently, His80^2.40^ toggles toward the receptor’s inner core and engages with Asp141^3.49^ in TM3, a feature that may help maintain the active conformation of the receptor (**Fig. 5H**).

### Abemaciclib inhibits ACKR3-mediated chemokine uptake *in vitro*

ACKR3 internalization (vide supra), previously linked to chemokine uptake, has been hypothesized to contribute to the shaping of chemokine gradients *in vivo* (12,14,50). Therefore, we next tested the impact of abemaciclib on the ACKR3-mediated chemokine uptake *in vitro* (**Fig. 6A**). Incubation of ACKR3-expressing HEK-293T cells, led to a rapid uptake of the ACKR3 ligands CXCL11-AF647 and CXCL12-AF647, but not fluorescent CCL5-AF647, whereas control cells showed no or very limited uptake of any of the chemokines (**Fig. 6C**). Co-incubation of ACKR3-expressing HEK-293T cells with 1 or 10 µM abemaciclib resulted in strongly reduced CXCL11-AF647 and CXCL12-AF647 uptake (**Fig. 6B and 6D**), in line with its ability to compete with the chemokines for ACKR3 binding.

**Figure 6:**
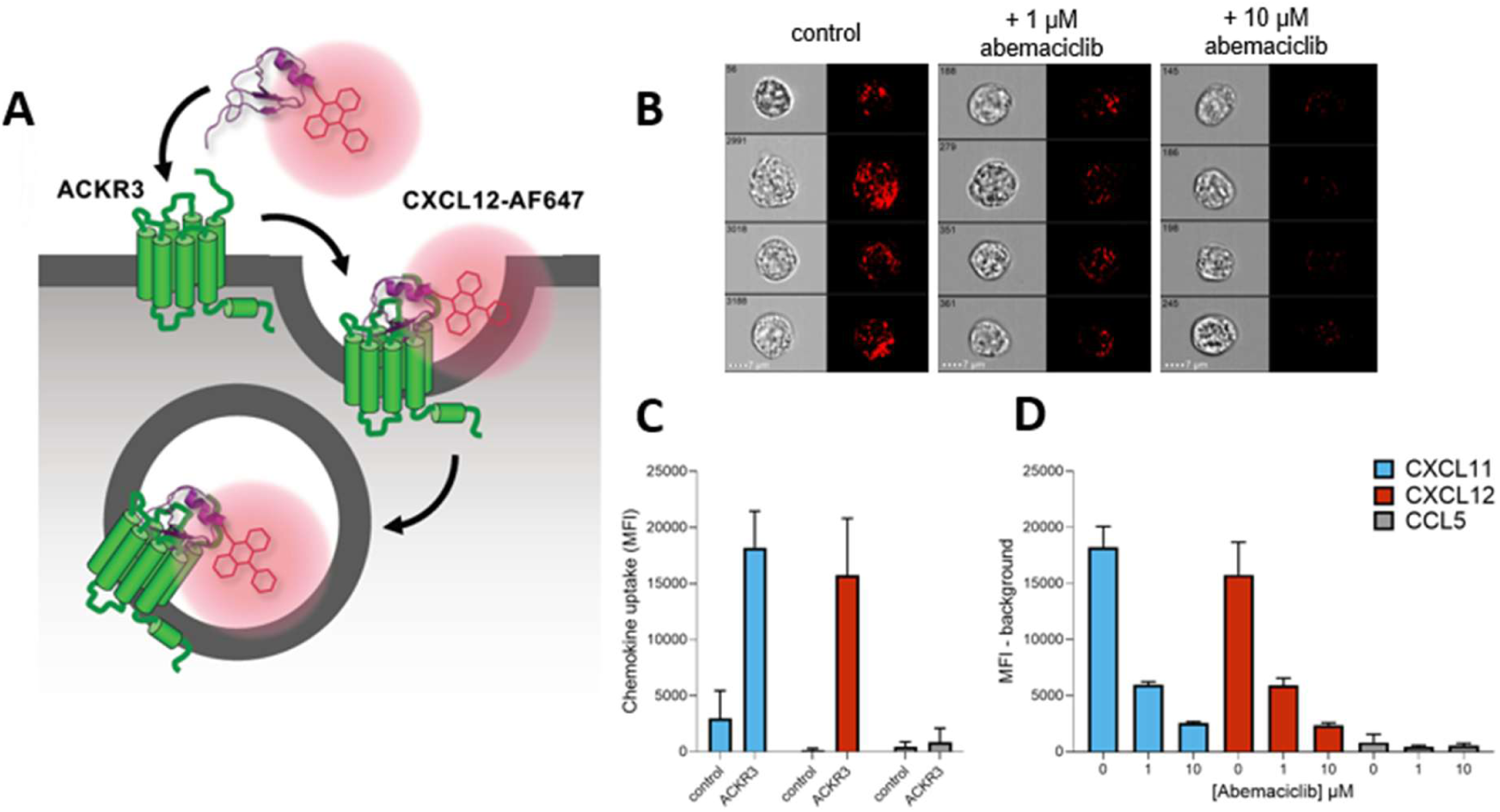
ACKR3-mediated CXCL11 and CXCL12 uptake is blocked by abemaciclib. A) Schematic representation of CXCL12-AZ647 internalization after binding to ACKR3. B) Four representative events of CXCL12-AF647 (2 nM, 60 min at 37°C) uptake with and without 1 or 10 µM abemaciclib pre-incubation in HEK-293T cells expressing ACKR3 visualized by imaging flow cytometry C) HEK-293T cells expressing ACKR3 take up more fluorescent CXCL11-AF647 and CXCL12-AF647, but not CCL5-AF647, compared to parental (control) HEK-293T cells, as quantified from imaging flow cytometry D) Abemaciclib prevents CXCL11 and CXCL12 uptake by HEK-293T cells expressing ACKR3 in a concentration-dependent manner, but has no effect on CCL5 uptake. Graphs represent pooled data from 3 individual experiments (error bars indicated respective S.D. values).

### *In vivo* ACKR3 target engagement by abemaciclib

Following up on the results the *in vitro* chemokine uptake assay, we tested the effect of abemaciclib on plasma CXCL12 levels of mice after abemaciclib exposure (**Fig. 7**). Previous studies have shown that plasma CXCL12 levels report *in vivo* target engagement of ACKR3 by small molecule drugs, like ACT-1004-1239 in mice and humans (28,51). Before the *in vivo* test, we evaluated if abemaciclib also interacts with mouse ACKR3 to rule out any potential species difference at the target level. As it can be seen in **Fig. 7B**, abemaciclib activates mouse ACKR3, as measured by NanoBiT-based β-arrestin-2 recruitment assay. Subsequently, mice were administered 75 mg/kg abemaciclib orally and plasma CXCL12 levels were measured 4 and 19 hours after administration. Whereas no difference in CXCL12 levels could be measured 4 hours after abemaciclib administration, a significant rise in plasma CXCL12 levels was observed after 19 hours (**Fig. 7C**).

**Figure 7:**
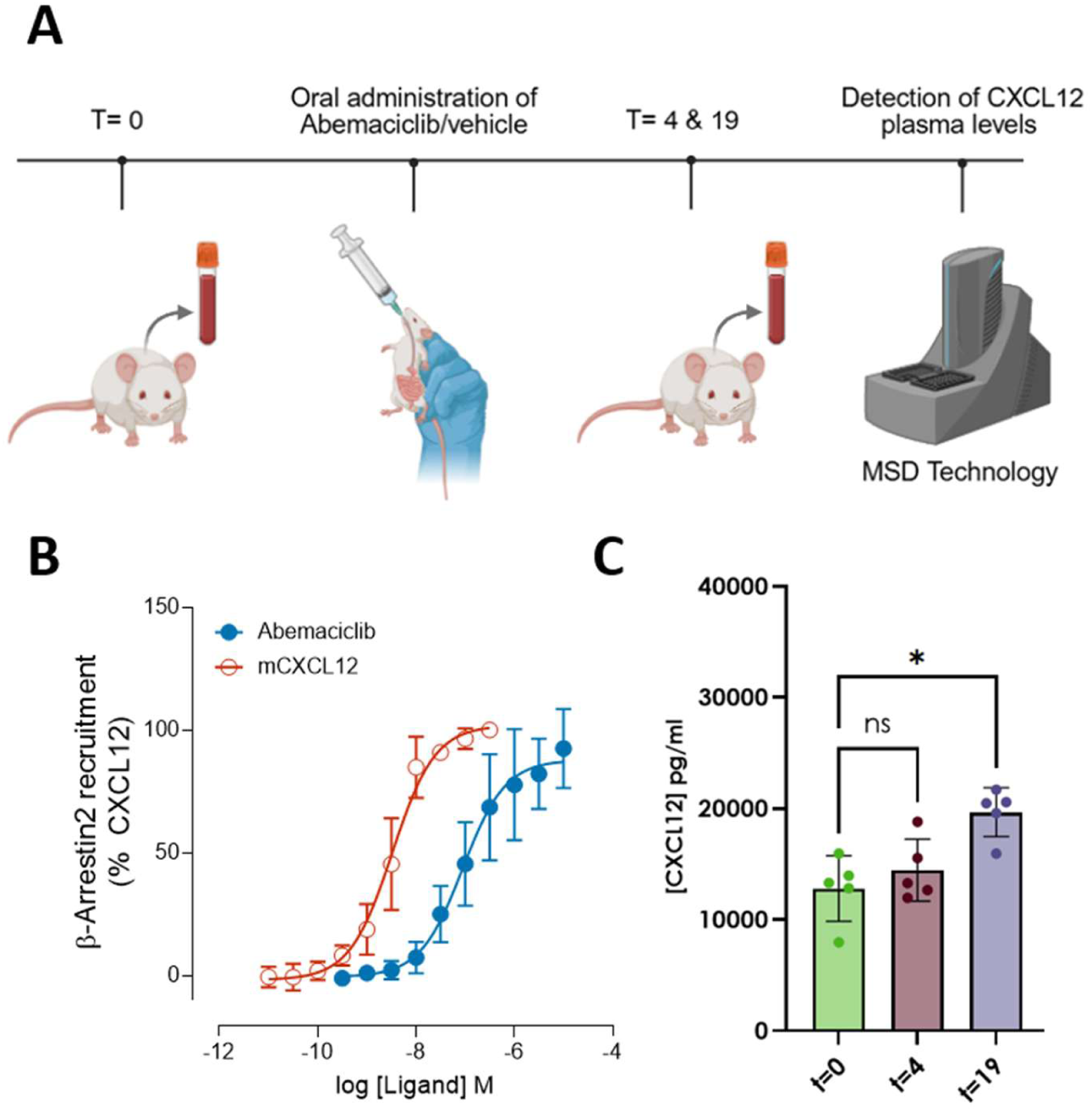
*In vivo* ACKR3 target engagement upon oral administration of abemaciclib. A) Schematic overview of mice administration of abemaciclib and quantification of CXCL12 in serum. B) HEK-293T cells were transfected with vectors encoding murine ACKR3-SmBiT and rat LgBiT-β-arrestin2 and stimulated for 15 minutes with abemaciclib or murine CXCL12. Recruitment of β-arrestin2 to mACKR3 was normalized to the mCXCL12 at the highest concentration tested (300 nM) Graphs represent pooled data from 4 independent experiments. Data points depicted are means, the error bars represent respective S.D values. C) Effect of a single oral administration (p.o.) of abemaciclib (75mg/kg) on CXCL12 serum concentrations in C57BL/6 mice. CXCL12 concentrations were measured 0, 4 and 19 hours after oral administration. Results are expressed as mean ± SD (n=5 per treatment group), using one-way analysis of variance (ANOVA) followed by Dunnett’s multiple comparisons test. *p <0.05.

## Discussion

The present study identifies the CDK4/6 inhibitor abemaciclib (Verzenios®) as a new structurally distinct modulator of ACKR3/CXCR7, thereby expanding the pharmacological landscape of this atypical chemokine receptor beyond the limited chemotypes that have been identified so far (30,31). Using a drug-repurposing strategy and a NanoBRET-based fluorescent chemokine binding assay (52), we found that abemaciclib binds ACKR3 with 25 nM affinity and displaces both CXCL12 and CXCL11 from the receptor, whereas other clinically approved CDK4/6 inhibitors fail to interact with ACKR3. The interaction is highly selective, as abemaciclib shows no measurable binding to CXCR4, ACKR5, or any of the other (atypical) chemokine receptors tested. Functionally, abemaciclib behaves as an ACKR3 agonist, inducing rapid β-arrestin-1 and 2 recruitment and promoting receptor internalization with potencies and efficacies comparable to the reference ACKR3 agonist VUF15485. Its major CYP3A4-derived metabolites retain ACKR3 binding and agonist activity, although with somewhat reduced potency. Following these findings, we capitalized on the recently developed experimental approaches in ACKR3 structural biology (10,46,47) to understand the molecular basis of ACKR3 binding and activation by this chemically distinct kinase inhibitor.

In the first ACKR3 cryo-EM study, two Fabs, CID24 and CID25, were used to determine structural data for ACKR3 bound to CXCL12 at 3.8 Å and the small-molecule agonist CCX662 at 3.8 Å resolution (47). These structures established that CXCL12 binds ACKR3 in a non-canonical orientation, rotated by ∼80° and shifted by ∼10 Å relative to canonical chemokine–receptor complexes, while its N-terminus still enters the orthosteric pocket (47). Subsequently, ACKR3 complexes with β-arrestin-1 or 2 were reported after GRK2- or GRK5-dependent phosphorylation, using Fab7 as an arrestin-stabilizing reagent (10). Recently, the structural landscape of ACKR3 was further extended by a series of active and inactive ACKR3 cryo-EM structures at 3.2–3.7 Å, including complexes with the agonists VUF11207, CXCL11, and VUF10661 and inverse agonists VUF16840 and VUN701 (48).

The cryo-EM structure of abemaciclib-bound ACKR3 reveals an active-like receptor conformation that aligns closely with the other agonist-bound ACKR3 structures. Interstingly however, abemaciclib binds differently from previously visualized ACKR3 agonists, wherein, it adopts an inverted U-shaped conformation and shows a relatively shallower binding site in the orthosteric pocket. This contrasts with the deeper binding of small-molecule agonists such as CCX662 (47) and VUF11207 (48), although its depth is closer to that of CXCL12-bound ACKR3 (47). The abemaciclib binding pose is stabilized by several specific interactions in the orthosteric pocket and extensive non-bonded contacts with residues contributed by TM1, TM2, TM3, TM6, TM7, and ECL1/ECL2, which is in agreement with its highly selective binding to ACKR3. Compared with the inactive VUF16840-bound state of ACKR3, abemaciclib induces hallmark rearrangements following GPCR activation (47,53,54), and therefore, further corroborates its ability to activate ACKR3 via converging molecular mechanism as observed for other well-established ACKR3 agonists.

NMR work on ACKR3 has provided an essential dynamic complement to the structural cryo-EM work by addressing how ligand binding reshapes receptor conformational ensembles rather than only stabilizing a single end state (47). Developing residue-specific ^13^CH_3_-methionine labelling of purified ACKR3 in micelles allowed solution NMR monitoring of conformational changes at native methionine residues distributed across the GPCR protein (46). Whereas Met212^5×39^ reports on the conformational changes in the extracellular/orthosteric ligand-binding region, Met138^3×46^, positioned near the NPxxY-associated intracellular region, reports on structural changes linked to effector coupling (46). Using ^13^CH_3_-methionine-labelled ACKR3, also abemaciclib binding was analysed by 2D ^1^H–^13^C HSQC NMR and compared with apo-ACKR3 and upon binding of chemokine CXCL12, small molecule agonist VUF15485, or inverse agonist VUF16840. Abemaciclib induces a clear chemical-shift perturbation of the Met138^3×46^ reporter relative to apo ACKR3, confirming that ligand binding is transmitted to the intracellular receptor core. Yet, the abemaciclib-induced Met138^3×46^ position does not overlap with the canonical reference positions defined by either VUF15485 or VUF16840, suggesting that abemaciclib stabilizes a distinct or more heterogeneous intracellular conformational ensemble than the conformational states of these reference ACKR3 small molecules.

Despite these differences in interaction and subsequent conformational ACKR3 dynamics, abemaciclib activates and internalizes ACKR3 protein with similar kinetics, potency and to the same extent as the reference agonist VUF15485. These findings underscore the present lack of understanding of how the observed conformational diversity of ACKR3 upon the binding of structurally different ligands, is linked to specific signal transduction pathways. The discovery of abemaciclib as a new ACKR3 ligand will further help to delineate these intricate details of ACKR3 function.

The presented *in vitro* and in *vivo* chemokine scavenging studies show the potential functional significance of ACKR3 agonism. Soon after the discovery of ACKR3 (then called CXCR7), chemokine scavenging by this “decoy” receptor was observed and presented to be involved in the generation of chemokine gradients and thereby e.g., modulate tumor microenvironment (12,55) or immune cell trafficking in Multiple sclerosis (MS) (21). Indeed, ACKR3 antagonism has been shown to increase plasma and tissue CXCL12 levels by preventing its scavenging, supporting CXCL12 elevation as a pharmacodynamic consequence of ACKR3 engagement (56). Similarly, abemaciclib could directly alter local CXCL12 availability in ACKR3-positive tumour, endothelial, or stromal compartments following ACKR3 binding.

While both small molecule ACKR3 antagonists and agonists functionally antagonize chemokine scavenging, their effect on ACKR3 expression might be different (30,31). Indeed, antagonists will not affect constitutive ACKR3 internalization and recycling, whereas inverse agonists increase ACKR3 surface expression (9,42). Oppositely, agonists increase ACKR3 internalization and might at least temporarily downregulate ACKR3 cell surface levels upon long-term exposure. ACKR3 is mostly recycled back to the cell membrane following CXCL12- and CXCL11-induced internalization, albeit with distinct kinetics (57), with very limited targeting for lysosomal degradation (7,58). Interestingly, increased ubiquitination-mediated proteasomal ACKR3 degradation was recently reported for the flavonoid derivative TBS6b, resulting in inhibition of hepatocellular carcinoma *in vitro* and *in vivo* (59). The long-term effect of abemaciclib exposure on cell surface ACKR3 availability remains to be investigated.

Clearly, with abemaciclib modulating chemokine scavenging *in vivo*, ACKR3 target engagement might be an important factor to fully understand its pharmacology. Abemaciclib is an orally available CDK4/6 inhibitor developed to exploit the cyclin D–CDK4/6–RB axis in hormone receptor-positive breast cancer (60–62). Mechanistically, it inhibits cyclin D-activated CDK4 and CDK6, thereby reducing retinoblastoma protein phosphorylation, maintaining E2F repression, and arresting cells in G1 phase (61). Clinically, CDK4/6 inhibitors are often grouped as one class, but the results of this study showed that they differ pharmacologically. Recently, abemaciclib was described as a “multi-functional radiation modifier,” building on preclinical data showing that abemaciclib enhances radiosensitivity, interferes with DNA-damage repair, and may affect radiation-induced vasculogenesis and antitumour immune responses (63,64). These effects are usually interpreted as consequences of CDK4/6 inhibition, but it is interesting to note that other ACKR3 ligands, such as the agonist CCX662, have been reported to improve anti-tumor responses after irradiation (65). Abemaciclib may therefore combine canonical CDK4/6-dependent radiosensitization with ACKR3-dependent modulation of the CXCL12 microenvironment. In ACKR3-rich tumours or irradiated vascular niches, abemaciclib could reshape CXCL12/CXCR4 signalling, potentially affecting tumour-cell migration, immune infiltration, endothelial repair, and vascular regrowth after radiotherapy.

In conclusion, our study identifies abemaciclib as a potent, structurally distinct ACKR3 agonist that shows a unique binding pose in the ACKR3 binding pocket, induces unique conformational ACKR3 dynamics and blocks *in vitro* and *in vivo* chemokine scavenging via ACKR3 engagement. These findings expand the ACKR3 ligand space, reveal an unexpected off-target mechanism for an important clinically used oncology drug, and provide a new scaffold for interrogating atypical chemokine receptor biology.

## Methods

### Pharmacological characterization of abemaciclib

#### Compounds and chemokines

Human recombinant CXCL8, CXCL11, CXCL12, CXCL13, CCL2, CCL5, CCL13, CCL19, CCL22, CCL25 and fluorescently labeled chemokines CXCL11-AZ594, CXCL11-AZ647, CXCL12-AZ488, CXCL12-AZ647, CCL5-AZ594, CCL5-AZ647 and CCL19-AZ594 were purchased from protein foundry (Milwaukee, WI, USA). Human recombinant CCL1, CCL3, CCL20, CCL27, CX3CL1 and CXCL16 were purchased from Peprotech, Thermo Fischer Scentific (Cranbury, NJ, USA). CXCL12-AF647 is purchased from Almac (Craigavon, UK). XCL2 was purchased from R&D systems (Minneapolis, MN, USA). Abemaciclib, Palbociclib and ribociclib and IT1t were purchased from Medchemexpress (Princeton, NJ, USA). Dalpiciclib was purchased from tebu-bio (Heerhugowaard, Netherlands). Trilaciclib, M2, M18 and M20 were purchased from GLPbio technology (Montclair, CA, USA). VUF15485 and VUF25550 were synthesized in-house. The FDA-approved drug library was purchased from TargetMol (Wellesley Hills, MA, USA).

#### Plasmids

Nluc-ACKR3 and Nluc-CXCR4 in pcDNA3.1 are kind gifts from prof S. Hill (Nottingham) (66). Nluc-ACKR2 is a kind gift of M. Szpakowska (67). Nluc-ACKR1 (GenBank ID: MN813503.1), Nluc-ACKR4 (GenBank ID: NP_057641.1) and Nluc-ACKR5 (GenBank ID: XM_054370942.1) were generated in a similar manner by substitution of ACKR3 and subcloned in pcDEF3 for Nluc-ACKR1 and Nluc-ACKR4. ACKR3-Smbit, Lgbit-β-arrestin1/2 and mVenus-CAAX have been previously described (9). pNBe vectors encoding GPCRs c-terminally tagged with SmBit and human β-arrestin-1 or β-arrestin-2 N-terminally fused to LgBit have been previously described (16).

#### Cell culture

Human Embryonic Kidney Cells (HEK293T cells) were maintained in DMEM supplemented with 10% fetal bovine serum (FBS), penicillin (100 µg/mL) and streptomycin (50 µg/mL) at 37°C, 5% CO2.

#### NanoBRET-based chemokine receptor binding assays on Membranes

Adherent HEK293T cells (4 million) were transiently transfected with 0.25 µg Nluc-ACKR1/pcDEF3, Nluc-ACKR3/pcDNA3.1, Nluc-ACKR5/pcDNA3.1, or Nluc-CXCR4/pcDNA3.1, or 0.5 µg with Nluc-ACKR2/pcDNA3.1 or Nluc-ACKR4/pcDEF3 using 30 µg of 25kDa linear polyethylenimine (L-PEI), as previously described (9). Total amount of plasmid DNA was kept at 5 µg per transfection using empty pcDEF3. After 48 hours, cells were harvested in ice-cold PBS, homogenised, ultracentrifuged and membranes were stored in 20 mM tris-sucrose at -80°C, pH = 7.4, as previously described (9). membranes were incubated with fluorescent chemokines (0.3 - 10 nM) and concentration series unlabeled ligands in white low-volume 384 well plates. with 10 nM CXCL11-AZ594, 0.3nM CXCL12-AF647, 10 nM CXCL12-AZ488, 10 nM CXCL12-AF647, 10 nM CCL5-AZ594 or CCL19-AZ594, respectively. All dilutions were prepared in HBSS (Thermo Fisher Scientific, Waltham, MA, USA) containing 0.2% BSA for Nluc-ACKR3 and Nluc-CXCR4, whereas Milli-Q-water (MQ) with 10 mM HEPES, 1 mM MgCl2, supplemented with 0.2% BSA was used for Nluc-ACKR1, Nluc-ACKR2, Nluc-ACKR4, and Nluc-ACKR5). After 1 hour incubation at 22°C, NanoGlo (1.4 µL/mL) (Promega, Madison, WI, USA) was added and BRET was measured after 5 minutes at wavelengths of the respective fluorophores (488, 594 or 647 nm) and nanoluc (460 nm) in the Pherastar plate reader (BMG Labtech, Ortenberg, Germany) at 22°C. Binding curves were analysed using a non-linear regression model with GraphPad Prism 10.2.3.

#### β-arrestin1/2 recruitment and ACKR3 internalization assays

HEK293T cells (1 million) were transfected in suspension with plasmids encoding for ACKR3-SmBit (0.4 µg), LgBit-β-arrestin1 (0.6 µg) or LgBit-β-arrestin2 (0.6 µg), pcDEF3 (1 µg) and 12 µg 25kDa L-PEI for the NanoBit-based β-arrestin2 recruitment assay, or with ACKR3-Nluc/pcDEF3 (0.05 µg), mVenus-CAAX/pcDEF3 (0.25 µg), pcDEF3 (1.7 µg) using 6 µg 25 kDa L-PEI for the NanoBRET-based ACKR3 internalization assay. Transfection mixes were incubated for 15 minutes at 22°C before adding to the cells. Subsequently 3 × 10^4^ cells/well were plated on a white 96 well plate and cultured for another 48 hours. On the assay day, the culture medium was replaced with HBSS supplemented with 0.05% BSA and cells were incubated for 15 minutes with ligand. Next, Nanoglo (1.4 µL/mL) was added, and bioluminescence was measured in the Pherastar plate reader at 37°C. For kinetic traces Nanoglo (1.4 µL/mL) was added first to measure baseline luminescence followed by the injection of ligands. Ligand-induced bioluminescence changes were calculated as fold change over vehicle-treated cells. Ligand-induced BRET changes for mVenus-CAAX internalization assay were measured at two wavelengths in the Pherastar plate reader and subsequently calculated by dividing the emission at 535-30 nm (mVenus) with emission at 475-30 nm (Nanoluc). Concentration response curves were analyzed by non-linear regression with GraphPad Prism 10.2.3 (GraphPad Software, San Diego, CA, USA).

#### Chemokine Uptake by Imaging Flow Cytometry

HEK293T, stably expressing ACKR3 were harvested in Opti-MEM supplemented with 1% BSA and distributed into 96-well plates (6 × 10^5^ cells/well). After pre-incubation for 10 min at 37°C with 0, 1, or 10 µM Abemaciclib MS, AZ647-labelled chemokines CXCL12, CXCL11, or CCL5 were added at a final concentration of 2 nM and the cells were incubated for 60 min at 37°C. Subsequently, the cells were washed with FACS buffer (1% BSA, 0.1% NaN3 in PBS) and Live/Dead stained with Zombie Green viability dye (Biolegend). After another washing step with FACS buffer, cells were treated with Cytofix fixation buffer (BD) for 20 min before a final wash with FACS buffer. Images of 1 × 10^4^ in-focus, living single cells were acquired with an ImageStream Mark II imaging flow cytometer (Amnis) equipped with an extended depth of field (EDF) module and using 60× magnification. Samples were analyzed using Ideas 6.2 software.

#### Chemokine receptor selectivity screen

HEK293T cells (1.8 × 10^6^ cells) were seeded in 6-well plates, and 24 h later, cotransfected with pNBE plasmids encoding chemokine receptors that are C-terminally fused to SmBiT and β-arrestin2 N-terminally fused to LgBiT, with the exception of CXCR4 that was C-terminally fused to LgBiT and combined with β-arrestin2 N-terminally fused to SmBiT, as previously described (16). After 24h, the cells were detached using Versene (0.48 mM EDTA in PBS) and resuspended in Opti-MEM and incubated for 25 min at 37°C with Nano-Glo Live Cell substrate and distributed into white 96-well plates (1 × 10^5^ cells/well). Cells were then treated with 10 µM abemaciclib or with 200 nM positive control agonist chemokine listed in the IUPHAR repository of chemokine receptor ligands. Ligand-induced β-arrestin2 recruitment to receptors was evaluated by recording bioluminescence signal for 20 minutes with a GloMax Discover plate reader (Promega). To evaluate antagonist properties of Abemaciclib, 20 nM of a full agonists of each receptor (100 nM chemokine for CCR9, CCR10 and XCR1) was added to the same wells and bioluminescence was measured again. Signal was normalized as percentage activity, where wells treated with a full agonist only were defined as 100% and signal from wells treated with no agonist was set to 0% activity.

### Cryo-EM structure of ACKR3-Abemaciclib

#### Expression and purification of ACKR3 for cryo-EM studies

The cDNA encoding full-length human ACKR3 was cloned in pVL1393 vector with N-terminus HA-signal, FLAG-tag followed by M4 sequence and HRV 3C protease site was expressed and purified from Spodoptera frugiperda (Sf9) insect cells using previously described protocols (68,69). Sf9 cells were infected with ACKR3-expressing baculovirus and harvested 72 h post-infection. Cell pellets were first homogenized in hypotonic buffer (20 mM HEPES pH 7.4, 20 mM KCl, 10 mM MgCl₂, 1 mM PMSF, 2 mM benzamidine) and subsequently extracted with hypertonic buffer (20 mM HEPES pH 7.4, 20 mM KCl, 10 mM MgCl₂, 1 M NaCl, 1 mM PMSF, 2 mM benzamidine).

For solubilization, cells were incubated in lysis buffer [20 mM HEPES pH 7.4, 450 mM NaCl, 1 mM PMSF, 2 mM benzamidine, 0.1% (w/v) cholesteryl hemisuccinate, 2 mM iodoacetamide] supplemented with 1% (w/v) L-MNG (Anatrace, Cat. no. NG310) for 2 h at 4°C. The lysate was then diluted with two volumes of dilution buffer (20 mM HEPES pH 7.4, 2.5 mM CaCl₂, 1 mM PMSF, 2 mM benzamidine) to adjust the NaCl concentration to 150 mM. After centrifugation at 47,936 × g for 30 min, the clarified supernatant was filtered and loaded onto pre-equilibrated M1-FLAG affinity resin.

The column was washed alternately with Low Salt Buffer (LSB; 20 mM HEPES pH 7.4, 150 mM NaCl, 2 mM CaCl₂, 0.01% cholesteryl hemisuccinate, 0.01% L-MNG) and High Salt Buffer (HSB; 20 mM HEPES pH 7.4, 350 mM NaCl, 2 mM CaCl₂, 0.01% L-MNG). The receptor was eluted using buffer containing 20 mM HEPES pH 7.4, 150 mM NaCl, 0.01% L-MNG, 2 mM EDTA, and 250 µg/mL FLAG peptide. To minimize receptor aggregation, free cysteine residues were alkylated by incubation with 2 mM iodoacetamide, followed by quenching with 2 mM L-cysteine. The purified ACKR3 receptors were supplemented with 10% (v/v) glycerol and stored at −80°C until further use.

#### Expression and Purification of CID24

CID24 was expressed and purified using a previously described protocol (70). The CID24 expression plasmid was transformed into E. coli BL21 (DE3) cells, and the primary culture was grown in 2×YT medium containing 100 μg/mL ampicillin at 37°C until an OD₆₀₀ of 1.0, at which point protein expression was induced with 1 mM IPTG. The culture was incubated for an additional 4 h at 37°C, after which the cells were harvested and resuspended in lysis buffer (20 mM HEPES pH 7.4, 500 mM NaCl, 0.5 mM MgCl₂, 1 mM PMSF, 2 mM benzamidine, 0.5% (v/v) Triton X-100).

Cells were lysed by sonication for 20 min at 40% amplitude, and the resulting lysate was heated at 65°C for 30 min. The sample was then centrifuged at ∼48,000 × g for 40 min at 4°C, and the clarified supernatant was filtered and loaded onto a pre-equilibrated Capto L (Cytiva) affinity column. The resin was washed with buffer containing 20 mM HEPES pH 7.4 and 500 mM NaCl, and the Fab fragment was eluted with 0.1 M acetic acid, followed by immediate neutralization using 10 times dilution of 1 M HEPES pH 8.0.

The purified protein was dialyzed against buffer containing 20 mM HEPES pH 7.4 and 150 mM NaCl, assessed by SDS–PAGE, flash-frozen in the presence of 10% (v/v) glycerol, and stored at −80°C until further use.

#### Reconstitution of ACKR3 complexes for structural analysis

ACKR3 complexes were assembled by incubating purified ACKR3 with a 1.3-fold molar excess of abemaciclib and approximately a 1.3-fold molar excess of CID24 for 1.5 h at room temperature. The resulting mixture was concentrated using a 100 kDa MWCO concentrator (Cytiva, Cat. no. 28932363) and subsequently loaded onto a Superose 6 Increase 10/300 GL column. Peak fractions corresponding to the desired complexes were assessed by SDS–PAGE, pooled, concentrated to 15–18 mg/mL, and used for cryo-EM grid preparation.

#### Cryo-EM grid preparation and data collection

Purified complex samples were applied onto glow-discharged Quantifoil R1.2/1.3 gold grids (300 mesh, holey carbon) at a protein concentration of 12 mg/ml (∼3 μl per grid). Grids were blotted for 4 s at 4°C and 100% relative humidity (blot force 0) using a Vitrobot Mark IV (Thermo Fisher Scientific) and immediately plunge-frozen in liquid ethane. Cryo-EM data were collected on a Titan Krios G4 electron microscope (Thermo Fisher Scientific) operated at 300 kV, equipped with a Gatan K3 direct electron detector and BioQuantum K3 energy filter. Automated data acquisition was performed using EPU software, with movies recorded in counting mode at a nominal magnification corresponding to a pixel size of 0.86 Å. Dose rates were maintained at ∼15.6 e⁻/Å²/s, with a total dose of 55 e⁻/Å² distributed over 50 frames for each 1 s exposure.

#### Cryo-EM data processing

The dataset for the abemaciclib+ACKR3+CID24 complex were processed following a standardized workflow in cryoSPARC v4.5-4.6, unless otherwise specified (71). Dose-fractionated movie stacks were corrected for beam-induced motion using the Patch motion correction (multi) sub-program, and contrast transfer function parameters were determined with Patch CTF. Particles were automatically picked using the blob-picker sub-program, applying a circular mask with a diameter range of 80-160 Å. The auto-picked particles were extracted with a box size of 300 px (Fourier cropped to 72 px) and subjected to reference-free 2D classification and heterogeneous refinement to eliminate ice contamination and distorted particles. Particles corresponding to the best 3D class were selected and re-extracted using a box size of 300 px (Fourier cropped to 256 px) and subjected to non-uniform (NU) refinement and local refinement. A complete data-processing pipeline including the number of particles selected at different stages are presented in Figure **S4**. Local resolution estimations for all cryo-EM maps were performed using the LocRes module in cryoSPARC, with half-maps provided as input. Protein-protein interfaces were analyzed using PDBsum1 (72).

#### Model building and refinement

Coordinates for ACKR3 were generated using SWISS-MODEL (73), and the atomic coordinates of CID24 were derived from the previously determined structure of the CCX662–ACKR3–CID24 complex (PDB: 7SK9) (47). All initial models were docked into the respective cryo-EM maps using UCSF Chimera, followed by flexible fitting using the “all-atom refine” function in COOT (74,75). Subsequent model refinement was performed with *phenix.real_space_refine*, applying secondary structure restraints, combined with iterative manual adjustments in COOT. The quality of the final models was assessed using MolProbity as implemented in Phenix (76,77). All structural figures were prepared with UCSF Chimera and ChimeraX (74).

### Nuclear Magnetic Resonance determination of ACKR3-abemaciclib complexes

#### ACKR3 protein expression and purification

Human full-length ACKR3 was expressed and purified as previously described (39). Briefly, baculovirus encoding ACKR3 was generated using the Bac-to-Bac Baculovirus Expression System (Invitrogen). The expression construct was assembled in a pFastBac1 plasmid with a gp64 promoter, an N-terminal HA signal sequence, a C-terminal PreScission protease cleavage site, and FLAG and 10xHis tags to aid in detection and purification. Sf9 cells were cultured at 27°C and 144 rpm in Expi293 media (Expression Systems). Large-scale expression of labeled ACKR3 was performed by adding high-titer baculovirus to a 2L culture of cells in methionine deficient medium (Expression Systems) at a density of 2.0-3.0 x 10^6^ cells ml^-1^ at an MOI of 5. After 5 hours, 250 mg/L ^13^CH_3_-methionine (Cambridge Isotope Laboratories) was added to cells. Cells were incubated at 27°C for 48 hours then pelleted and stored at -80°C prior to purification.

ACKR3 was purified as previously described (39). Briefly, cells containing 13CH3-methionine ACKR3 were lysed by needle/syringe before the addition of ligand (2 uM abemaciclib) for initial complex formation for 30 min. Solubilization buffer (100 mM HEPES pH 7.5, 800 mM NaCl, 1.5% / 0.3% MNG/CHS) was added and spun for 4 hours at 4°C. After solubilization, slurry was spun down at 50,000x g for 30 min to isolate solubilized receptor. Solubilized receptor was transferred to conical tubes and rotated overnight in the presence of TALON cobalt resin with 10 mM imidazole. Conical tubes were spun down and resin transferred to gravity column for further purification. Resin was washed with Wash Buffer 1 (50 mM HEPES pH 7.5, 400 mM NaCl, 0.1%/0.02% MNG/CHS, 10% glycerol, 20 mM imidazole), Wash Buffer 2 (50 mM HEPES pH 7.5, 400 mM NaCl, 0.025%/0.005% MNG/CHS, 10% glycerol, 10 mM imidazole), and eluted with Elution Buffer (50 mM HEPES pH 7.5, 400 mM NaCl, 0.025%/0.005% MNG/CHS, 10% glycerol, 250 mM imidazole). Eluted ACKR3 was concentrated down to 500 uL and exchanged into Exchange Buffer (25 mM HEPES pH 7.5, 150 mM NaCl, and 0.025%/0.005% MNG/CHS)) using a PD-10 desalting column (GE). Cleavage of the His-FLAG purification tag and N-linked glycosylations was performed overnight at 4°C with PreScission Protease and PNGaseF, respectively. Following tag cleavage, ACKR3 was applied to a reverse TALON column, and flow-through was isolated and concentrated down to ∼300 uL for NMR analysis.

#### ACKR3 Nuclear Magnetic Resonance (NMR) measurements

Heteronuclear single quantum coherence (HSQC) spectra were collected from purified ACKR3 samples in 10% D_2_O loaded into a 5 mm Shigemi microtube on a Bruker Avance 800 MHz spectrometer with a triple resonance cryogenic probe at 310K for 27 hr. Data were processed using NMRPipe (78) and plotted in XEASY (79).

### CXCL12 quantification in mice serum

Female C57BL/6 mice (8 weeks old) were purchased from Janvier-Labs. All experiments involving laboratory animals were conducted in a pathogen-free animal facility at the Luxembourg Institute of Health with the approval of the Luxembourg Ministry for Agriculture (#LUPA 2021/22). Mice were treated following the European guidelines.

Abemaciclib was formulated in 1% hydroxyethyl cellulose (Merck, Darmstadt, Germany) in 25 mM phosphate buffer (pH 2.0). A single dose of abemaciclib (75 mg/kg) or vehicle (1% hydroxyethyl cellulose in 25 mM phosphate buffer) was administered by oral gavage. Serum samples were collected before (t=0) or 4 and 19 hours after administration.

The CXCL12 concentration in serum samples was measured using the U-PLEX Mouse SDF-1α Assay (kit: K152VBK-1; Meso Scale Discovery, Rockville, MD, USA) according to the manufacturer’s instructions. Data acquisition was conducted using the MESO Quickplex SQ 120 instrument, and the subsequent analysis was performed using the MSD Discovery Workbench 4.0.13 software (LSR_4_0_13).

## Data availability

The cryo-EM structure of abemaciclib-CXCR7 and corresponding atomic coordinates have been deposited in the Electron Microscopy Data Bank (EMDB) and Protein Data Bank (PDB) under accession codes; EMD-81385 and pdb_000027sf, respectively.

## Acknowledgments

The research on chemokine receptors in the A.K.S. laboratory is supported by the Department of Biotechnology (BT/PR53019/MED/30/2528/20), J. C. Bose Grant of ANRF (ANRF/JBG/2025/000055/LS), and the Lady Tata Memorial Trust. The cryo-EM structure of abemaciclib-CXCR7 was determined at the ANRF-Sponsored National cryo-EM Facility at IIT Kanpur supported by ANRF/SERB (IPA/2020/000405). We thank Ravi Ranjan and Pradeep Gautam for assistance with cryo-EM grid freezing and data collection. NMR spectra were acquired on an 800 MHz instrument supported by NIH grant S10 OD25000. B.F.V. and S.J. were supported by NIH grant R37 AI058072. This study was supported by the Luxembourg Institute of Health (LIH) through the NanoLux Platform, the Cancer Foundation Luxembourg, Luxembourg National Research Fund (INTER/FNRS CXCL12 20/15084569, CORE IMPACTT C23/BM/18068832, ACKROS C25/BM/19563724, INTER/AUDACE/25/19352183, PRIDE 21/16763386/CANBIO2).

## Author Contributions

M.C.M.L.B., J.D., D.N., M.M., S.J., N.B., M.G., A.R., R.B., E.A.L., Y.K., and J.P. designed and performed experiments and analysed the data. S.Z., D.P., and A.H., contributed to study conceptualization and manuscript review. B.F.V, H.F.V., A.K.S., A.C., M.S., and R.L supervised and coordinated the research. M.C.M.L.B., H.F.V., and R.L. wrote the original manuscript, with contributions from J.D., D.N., S.J., N.B. and M.M. All authors reviewed and edited the manuscript.

## Notes

### Competing Interest Statement

The authors have declared no competing interest.

