## Supplemental Figures for "The FDA-approved CDK4/6 inhibitor Abemaciclib is a potent agonist of the β-arrestin-biased atypical chemokine receptor ACKR3"

### Supplementary information

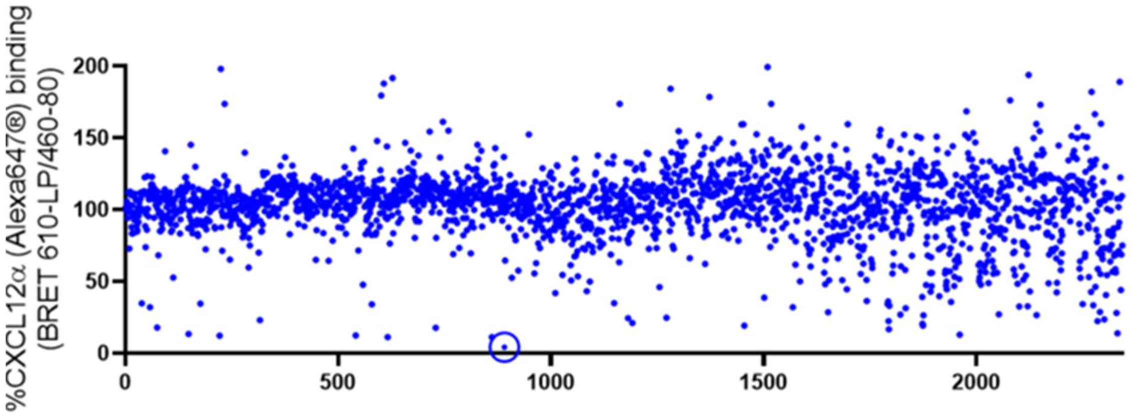


**Figure S1: FDA approved drug library screen against ACKR3**

10 µM of a library consisting of 2342 FDA approved drugs were screened for displacement of 0.3 nM CXCL12-AF647 on membranes from HEK-293T cells expressing NLuc-ACKR3. Circled in blue is abemaciclib. Data depicted is a single experiment in monoplo.

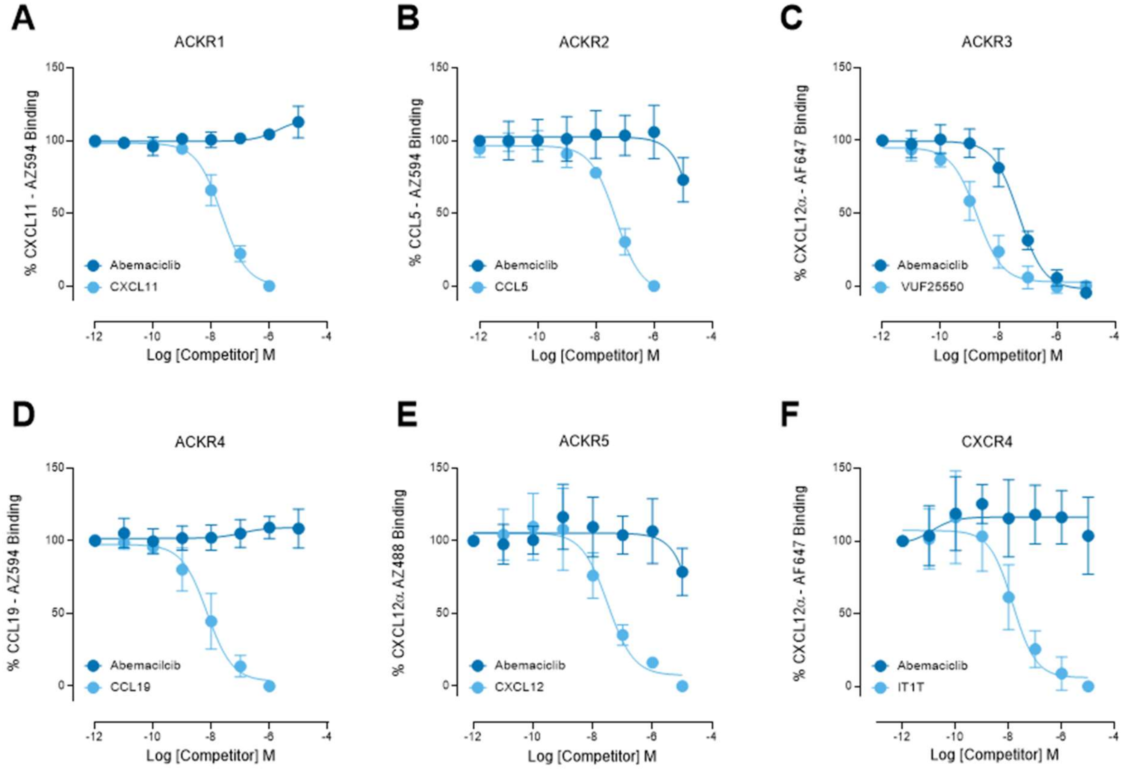


**Figure S2: Abemaciclib selectively binds to ACKR3**

Membranes of HEK-293T cells transiently expressing either A) NLuc-ACKR1, B) NLuc-ACKR2, C) NLuc-ACKR3, D) NLuc-ACKR4, E) NLuc-ACKR5 or F) NLuc-CXCR4 were used to measure the concentration dependent displacement of binding of 10 nM CXCL11-AZ594, 10 nM CCL5-AZ594, 0.3 nM CXCL12-AF647, 10 nM CCL19-AZ594, 10 nM CXCL12-AZ488, or 10 nM CXCL12-AF647, respectively. Percentage of remaining fluorescently labelled chemokine was normalized. Graphs represent pooled data from 3 independent experiments. Data points depicted are means, the error bars represent respective S.D. values.

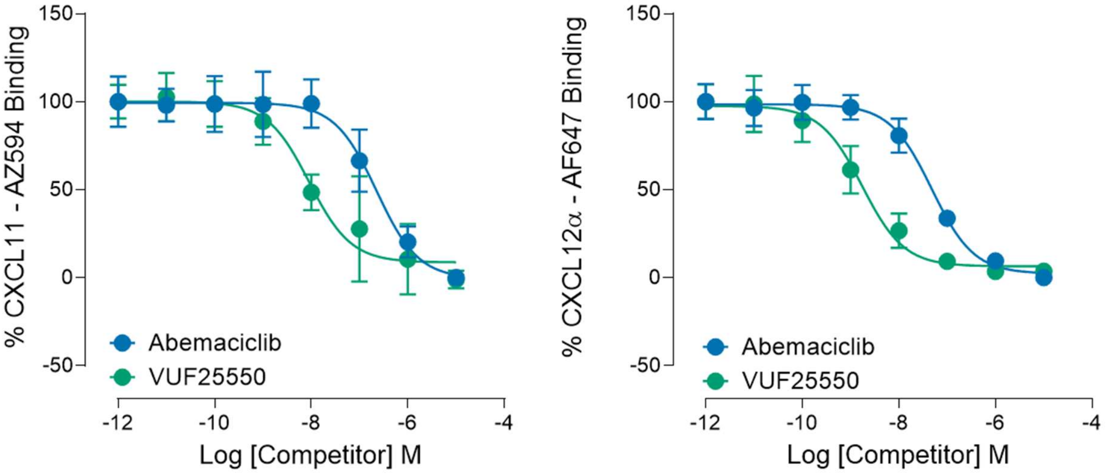


**Figure S3: Abemaciclib displaces both CXCL11 and CXCL12 from NLuc-ACKR3**

Membranes of HEK-293T cells transiently expressing NLuc-ACKR3 were used to measure the concentration dependent displacement of binding of A) 0.3 nM CXCL11-AZ594 and B) 0.3 nM CXCL12-AF647 by abemaciclib and inverse agonist VUF25550. Percentage of remaining fluorescently labelled chemokine was normalized. Graphs represent pooled data from 3 independent experiments. Data points depicted are means, the error bars represent respective S.D. values.

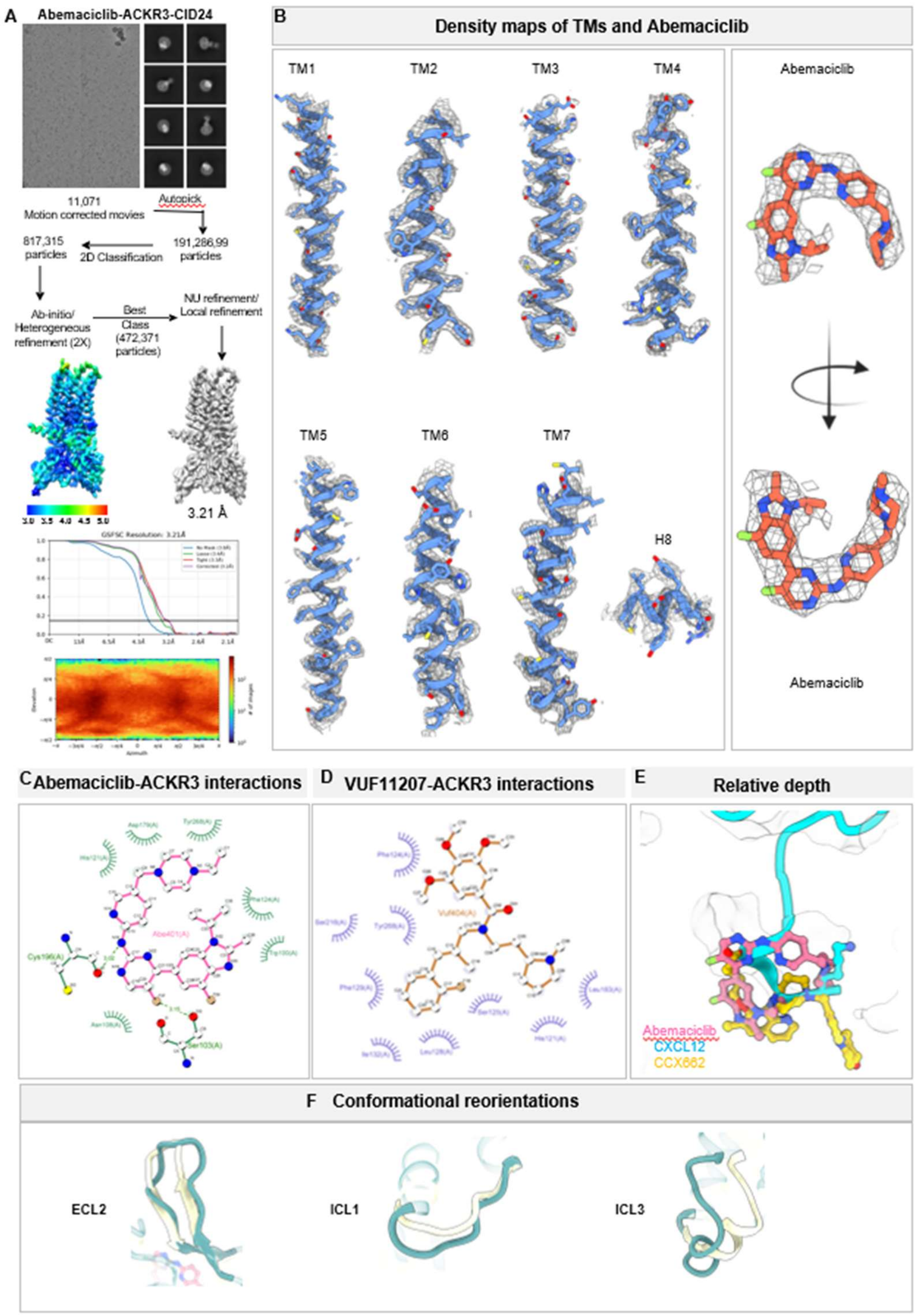


**Figure S4: Structure determination and ligand-binding interactions of ACKR3.**

A) Schematic representation of the data processing pipeline followed for the structure determination of the abemaciclib-ACKR3 dataset. B) Representative densities and fitted models of TM1-TM7 and abemaciclib are shown. C, D) Schematic representation illustrating the binding interactions of abemaciclib (C) and VUF11207 (D) with ACKR3, respectively. E) Ball and stick representation of relative depth of binding of abemaciclib, CXCL12 variant, and CCX662 within the orthosteric pocket of ACKR3. F) Superimposition of agonist abemaciclib- and inverse agonist VUF16840-bound ACKR3 structures highlighting conformational reorientations observed in ICL1, ICL3, and ECL2.
